# Heat dissipation drives the hump-shaped scaling of animal dispersal speed with body mass

**DOI:** 10.1101/2022.09.08.507078

**Authors:** Alexander Dyer, Ulrich Brose, Emilio Berti, Benjamin Rosenbaum, Myriam R. Hirt

**Affiliations:** EcoNetLab, German Centre for Integrative Biodiversity Research Halle-Jena-Leipzig, Leipzig, Germany; Institute of Biodiversity, Friedrich-Schiller-University Jena, Jena, Germany

## Abstract

Dispersal is critical to animal survival and thus biodiversity in fragmented landscapes. Increasing fragmentation in the Anthropocene necessitates predictions about the dispersal capabilities of the many species that inhabit natural ecosystems. This requires mechanistic, trait-based models of animal dispersal which are sufficiently general as well as biologically realistic. While larger animals should generally be able to travel greater distances, reported trends in their speeds across a range of body sizes suggest limited locomotor capacities among the largest species. Here, we show that this also applies to dispersal speeds and that this arises because of their limited heat-dissipation capacities. We derive a model considering how fundamental biophysical constraints of animal body mass associated with energy utilisation (i.e. larger animals have a lower metabolic energy cost of locomotion) and heat-dissipation (i.e. larger animals require more time to dissipate metabolic heat) limit sustained (i.e. aerobic) dispersal speeds. Using an extensive empirical dataset of animal dispersal speeds (531 species), we show that this *allometric heat-dissipation model* best captures the hump-shaped trends in dispersal speed with body mass for flying, running and swimming animals. This implies that the inability to dissipate metabolic heat leads to the saturation and eventual decrease in dispersal speed with increasing body mass as larger animals must reduce their realised dispersal speeds in order to avoid hyperthermia during extended dispersal bouts. As a result, the highest dispersal speeds are achieved by animals of intermediate body mass, whereas the largest species might suffer from stronger dispersal limitations in fragmented landscapes than previously anticipated. Consequently, we provide a mechanistic understanding of animal dispersal speed that can be generalised across species, even when the details of an individual species’ biology are unknown, to facilitate more realistic predictions of biodiversity dynamics in fragmented landscapes.

## Introduction

Dispersal is a fundamental process in ecology and evolution that drives biodiversity patterns by linking local populations to one another across several temporal and spatial scales [1,2]. Actively dispersing animals are capable of colonising distant habitat patches that provide access to novel resources, predator-free environments, and reproductive opportunities. This has substantial implications for their reproduction and survival. Despite these potential benefits, however, dispersal carries with it considerable costs. For instance, during transience - the phase of dispersal which enables these large-scale displacements [3,4] - animals must contend with extended bouts of elevated metabolic expenditure in order to sustain locomotion through a, typically, hostile landscape (reviewed in [5]). Such trade-offs contribute to the emergence of complex spatial dynamics within landscapes (e.g. allee-, rescue-, source-sink, and drainage effects) that are crucial to species persistence [6-**Error! Reference source not found**.8]. In light of mounting evidence suggesting that anthropogenic perturbations such as landscape fragmentation, climate change, and land-use change disrupt the natural movements of animals [**Error! Reference source not found**.-**Error! Reference source not found**.**Error! Reference source not found**.**Error! Reference source not found**.], a general mechanistic understanding of animals’ dispersal capacities represents a critical step towards addressing the consequences of landscape connectivity for patterns of biological diversity [**Error! Reference source not found**.-**Error! Reference source not found**.17].

A central component of an animal’s dispersal capacity is its sustained (i.e. aerobic) speed, which fundamentally depends on its locomotion mode, body mass and the temperature which it experiences while dispersing. Unlike the speeds of foraging animals, which can be predicted by optimal foraging theory, the speeds of dispersing animals depend on several context-dependent trade-offs between energy costs, travel time, and dispersal distance [18,**Error! Reference source not found**.]. Among animals of similar size, flying is generally faster than running and swimming, while within each locomotion mode, larger animals usually move faster and farther than smaller animals ([20-22], but see [**Error! Reference source not found**.,**Error! Reference source not found**.]).

Allometric relationships have achieved generality in relating animal body mass to the mechanical-[25-**Error! Reference source not found**.**Error! Reference source not found**.**Error! Reference source not found**.] and metabolic energy costs of locomotion [**Error! Reference source not found**.-**Error! Reference source not found**.**Error! Reference source not found**.32]; however, devising a general allometric model that can predict the speeds that animals are capable of sustaining has remained a challenge: Among existing models which, typically, describe a power-law scaling relationship between speed and animal body mass (but see [33]), there are significant discrepancies in the reported values of the allometric scaling exponent across disparate groups of flying, running and swimming animals (e.g. [21,34-**Error! Reference source not found**.**Error! Reference source not found**.38]). This suggests that models based solely on the mechanical- and metabolic energy demands of locomotion are insufficient to predict the sustained speeds of animals across a sufficiently wide range of taxonomic groups and locomotion modes.

Based on prior models of maximum speed [**Error! Reference source not found**.,40], we also consider a hump-shaped relationship between sustained speed and body mass. So far, very few studies have explicitly considered the importance of metabolic heat - an inevitable by-product of muscular contractions - in limiting the speeds that animals can sustain during extended locomotion bouts (but see [41-**Error! Reference source not found**.**Error! Reference source not found**.]). In order for an animal’s core body temperature to remain stable and within its thermal limits, it is essential that the heat that its body dissipates to the ambient environment is sufficient to balance the excess heat that its muscles produce during locomotion. If heat-dissipation cannot offset metabolic heat production, animals must decrease their metabolic demands and, thus, their speed in order to avoid hyperthermia. Consequently, we argue that a general mechanistic model that can predict the sustained speeds at which animals can actively disperse must account for both the fate of energy that goes towards the performance of useful work as well as the fate of energy that is dissipated internally as heat.

We derive a general allometric model that considers how fundamental biophysical constraints associated with the supply, utilisation and dissipation of energy and heat limit the sustained speeds of flying, running, and swimming animals during their transient phase of dispersal. Our model builds on previous biomechanical and metabolic approaches by considering the body-mass dependence of (1) aerobic metabolism [44,45] and (2) the metabolic cost of locomotion [**Error! Reference source not found**.-**Error! Reference source not found**.**Error! Reference source not found**.]. Unlike previous models, we also consider how (3) animals’ capacity to dissipate metabolic heat fundamentally constrains their capacity for sustained locomotion. This ultimately leads to the prediction of a hump-shaped relationship between sustained speed and body mass. We established an exhaustive dataset on empirical animal dispersal speeds to test (1) whether this new heat-dissipation model provides more accurate predictions of animal dispersal speeds than conventional power-law models and (2) if it makes consistent predictions across locomotion modes and ecosystem types. Ultimately, our approach provides a mechanistic model of animal dispersal speed that can be generalised across species, even when the details of an individual species’ biology are unknown.

## Model development

We derive three alternative models of how animal dispersal speed scales with body mass. The models are based on varying assumptions of how, for a given distance moved, the total time budget that is allocated towards dispersal is split into moving time and heat-dissipation time (Fig. 1a). For simplicity, we retain the concept of discrete time budgets for movement and heat-dissipation, while empirically both can take place at infinitely small time-steps (e.g. between each stride) without violating the assumptions of the concept (Fig. 1a, lowest bar). In other words, animals do not need to stop to dissipate heat; instead, they continuously allocate part of their total time budget towards heat-dissipation by moving more slowly. The three models are based on the same allometric relationships for metabolic power generation and locomotion efficiency and therefore predict the same potential dispersal speed (Fig. 1b). However, they differ in whether they assume that heat-dissipation time is (1) not necessary (*metabolic model*), (2) constant across all species (*constant heat-dissipation model*), or (3) increases with body mass (*allometric heat-dissipation model*, Fig. 1c). Consequently, they predict that the realised dispersal speed scales with body mass as a power law (*metabolic model*), a saturating function (*constant heat-dissipation model*), or a hump-shaped function (*allometric heat-dissipation model*, Fig. 1d). In the following, we provide an overview of the model derivation (see also Table 1), while the detailed derivation is provided as a supplement (see S1 Text).

**Table 1.** Overview of three alternative allometric locomotion models and their corresponding mechanistic hypotheses.

|  | Metabolic model | Constant heat-dissipation model | Allometric heat-dissipation model |
| --- | --- | --- | --- |
| Step 1: Dispersal speed | $v = \frac{D}{t_{total}}$ | | |
| Step 2: Distance moved | $D = P_i \cdot E_{max} \cdot t_{move}$ | | |
| Step 3: Metabolic power input | $P_i = P_0 \cdot M^a$ | | |
| Step 4: Maximum locomotion efficiency | $E_{max} = E_0 \cdot M^b$ | | |
| Step 5: Potential dispersal speed | $v = \frac{(P_0 \cdot E_0) \cdot M^{a+b} \cdot t_{move}}{t_{total}} = \frac{v_0 \cdot M^c \cdot t_{move}}{t_{total}}$ | | |
| Step 6: Total dispersal time, movement time and heat-dissipation time | $t_{total} = t_{move}$ | $t_{total} = t_{move} + t_{diss}$ | |
| Step 7: Scaling of heat-dissipation time | $t_{diss} = 0$ | $t_{diss} = k_0 \cdot D$ | $t_{diss} = k_\lambda \cdot M^d \cdot D$ |
| Step 8: Full models | $v = v_0 \cdot M^c$ | $v = \frac{\frac{1}{k_0} \cdot M^c}{M^c + \frac{1}{v_0 \cdot k_0}}$ | $v = \frac{\frac{1}{k_\lambda} \cdot M^c}{M^{c+d} + \frac{1}{v_0 \cdot k_\lambda}}$ |

Description of model parameters: dispersal speed, $v$ ( $m s^{-1}$ ); dispersal distance, $D$ ( $m$ ); total dispersal time, $t_{move}$ ( $s$ ); animal body mass, $M$ ( $kg$ ); locomotion rate constant that varies among locomotion modes, $v_0$ ( $m s^{-1} kg^{-c}$ ); heat-dissipation time constants, $k_0$ ( $s m^{-1}$ ) and $k_\lambda$ ( $s m^{-1} kg^{-d}$ ); scaling exponents describing the body mass-dependence of metabolic power input, $a$ , maximum locomotion efficiency, $b$ , potential dispersal speed, $c$ , and heat-dissipation time, $d$ .

**Fig 1.**
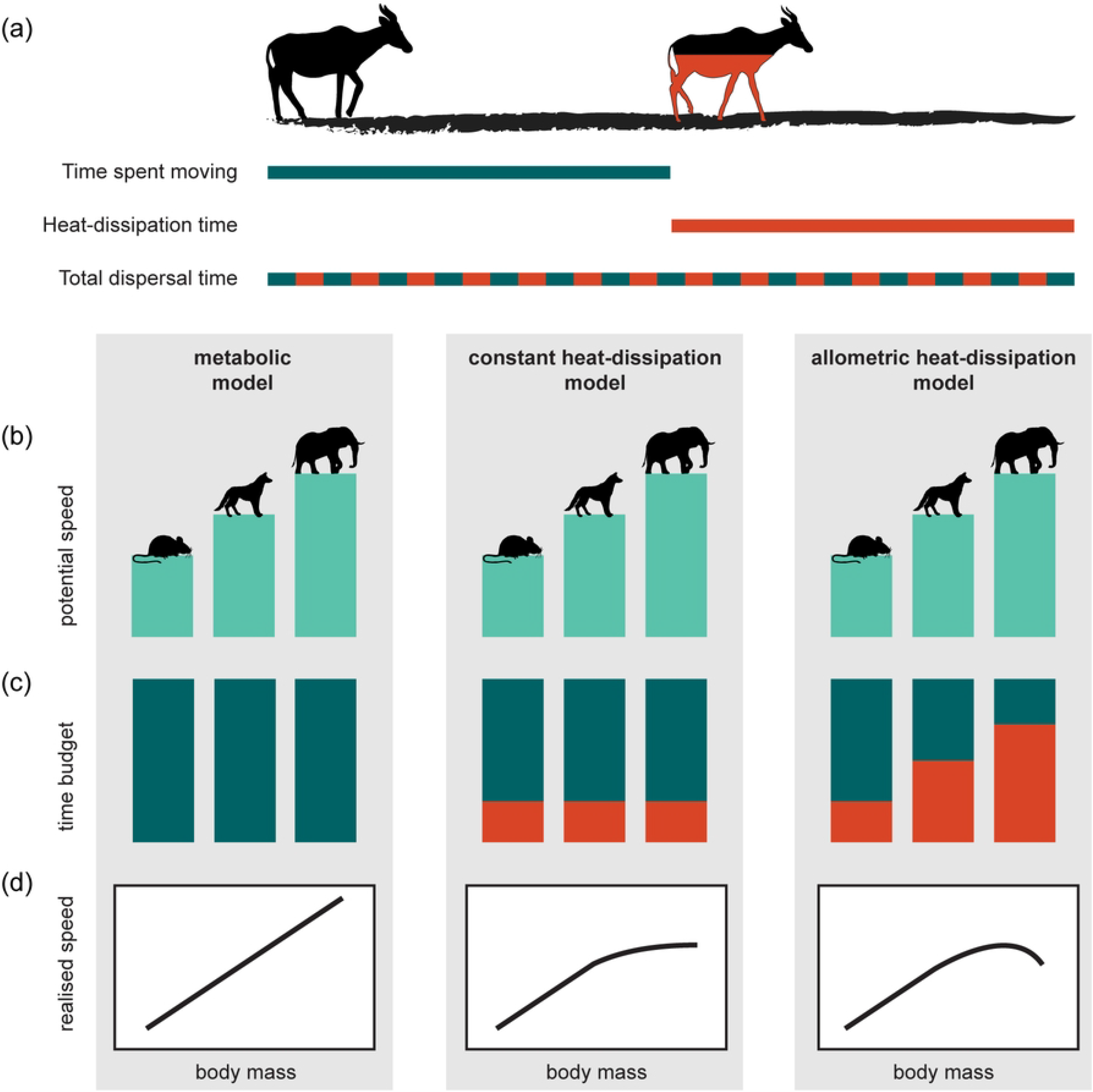
Concept of heat-dissipation time as a fundamental constraint to the realised speeds of dispersing animals. (a) Dispersal speed represents the distance dispersed divided by the total time allocated towards dispersal. The time spent moving a unit distance is associated with the production of metabolic heat by contracting muscles as they perform the mechanical work required for locomotion. In order for the core body temperature to remain stable, a fraction of the total dispersal time is allocated towards heat-dissipation to offset the heat that is produced during locomotion; this takes place at small time-steps (e.g. between each stride). An increase in heat-dissipation time, therefore, corresponds to a reduced stride frequency and a net decrease in the realised dispersal speed. (b) A greater supply of metabolic power combined with a higher locomotion efficiency allows larger animals to sustain higher potential dispersal speeds. (c) The fraction of the total dispersal time budget that is allocated towards locomotion (blue) or heat-dissipation (red): (1) time is exclusively allocated towards locomotion (*metabolic model*); (2) all species allocate a constant (i.e. body mass-independent) fraction towards heat-dissipation (*constant heat-dissipation model*); (3) larger animals allocate a larger fraction towards heat-dissipation (*allometric heat-dissipation model*). (d) The allocation of heat-dissipation time determines the realised dispersal speed that can be sustained, yielding a (1) power-law (*metabolic model*), (2) saturating (*constant heat-dissipation model*), or (3) hump-shaped (*allometric heat-dissipation model*) allometric scaling model.

All three allometric models of dispersal speed have the first five steps of model derivation in common: First, dispersal speed is equal to the distance moved divided by the total dispersal time (Table 1, step 1). Second, distance moved is predicted by the product of whole-organism metabolic power input and locomotion efficiency (Table 1, step 2). Third, metabolic power input scales with body mass as a power law (Table 1, step 3). Fourth, the maximum locomotion efficiency, the reciprocal of the minimum absolute metabolic cost of locomotion, also follows a power-law scaling relationship with body mass (Table 1, step 4). Taken together, these terms produce an allometric model of potential dispersal speed that is shared by all three models (Table 1, step 5, Fig. 1B). The three models differ in their assumptions on time budgets (Table 1, step 6, Fig. 1C) and the body-mass constraints associated with heat-dissipation time (Table 1, step 7). The simple *metabolic model* implicitly assumes that all animals dedicate their total dispersal time, *t*_*total*_ (*s*), exclusively towards sustained locomotion and therefore disperse at speeds that minimise their absolute metabolic cost of locomotion. This yields a power-law scaling of dispersal speed with body mass (Table 1, step 8 first column, Fig. 1D). Both heat-dissipation models assume that the total dispersal time is not entirely dedicated towards locomotion as dispersing animals allocate some time, *t*_*diss*_ (*s*), towards the dissipation of metabolic heat - an inevitable by-product of muscular contractions (Table 1, step 6). Rather than accelerating and decelerating from rest, we assume that animals allocate heat-dissipation time at small time-steps throughout the locomotion process, for instance, between each stride (conceptualised in Fig. 1A, lowest bar). An increase in heat-dissipation time therefore corresponds to a reduced stride frequency and a decrease in the realised speed that can be sustained. Our second model, the *constant heat-dissipation model*, is a saturating (non-decreasing) allometric scaling model (Fig. 1D) which assumes, implicitly, that all animals possess the necessary physiological and/or behavioural adaptations to adequately facilitate the dissipation of metabolic heat via body mass-independent pathways (Table 1, steps 7-8 middle column). Our final model, the *allometric heat-dissipation model*, is a hump-shaped allometric scaling model (Fig. 1D). It includes the additional assumption that the maximum heat-dissipation capacity of animals, and thus, the additional time that must be allocated towards heat dissipation, also scales with body mass (Table 1, step 7 right column). This implies that larger animals have greater thermal inertia - their body temperature changes more slowly than that of smaller animals when experiencing the same thermal gradient. Accordingly, larger animals require more time to dissipate the heat that is produced while moving a unit distance. A consequence of this body mass-dependent allocation of additional heat-dissipation time is that the largest animals must experience a net reduction in their sustained speed in order to effectively regulate their body temperature during the extended locomotion bouts. This yields a hump-shaped scaling of realised dispersal speed with body mass (Table 1, step 8 right column, Fig. 1D).

## Results

We compared the ability of three hypothesis-driven models (see Table 1) to predict the dispersal speeds of animals across three different modes of locomotion. Model comparison using LOOIC showed that the *allometric heat-dissipation model* (Table 1 step 8) best describes the systematic relationship between body mass and realised dispersal speed across flying, running, and swimming animals while the *metabolic model* and the *constant heat-dissipation model* score substantially worse (Table 2).

**Table 2.** Comparison of three alternative allometric locomotion models that predict the realised dispersal speeds of animals as a function of their body mass and locomotion mode. LOOIC values are presented together with the difference in LOOIC value relative to the most parsimonious model (ΔLOOIC = 0.0) and the estimated standard error of the difference (SE ΔLOOIC). All models fit the locomotion rate constant, *v*_0_, independently (i.e. no pooling) for flying, running, and swimming animals.

| Model | Description | LOOIC | $\Delta\text{LOOIC}$ | SE $\Delta\text{LOOIC}$ |
| --- | --- | --- | --- | --- |
| Metabolic | Power law | 465.0 | 62.8 | 14.4 |
| Constant heat-dissipation | Saturating | 427.6 | 25.4 | 5.8 |
| Allometric heat-dissipation | Hump-shaped | 402.2 | 0.0 | 0.0 |

After accounting for differences in the locomotion rate constant, *v*_0_, among flying, running and swimming animals, the *allometric heat-dissipation model* predicts that a single hump-shaped relationship (in log-log space) can be used to describe the realised dispersal speeds of all species as a function of their body mass (Fig 2; Table 3). We also tested a slightly more complex *allometric heat-dissipation model* that accounts for variation in the allometric scaling exponent *c* across the three locomotion modes (S3 Fig). This more complex model yielded comparable prediction accuracies to that of the best-performing model (ΔLOOIC = 2.0, SE ΔLOOIC = 5.3, S1 Table). We have, therefore, focused on the results of the more parsimonious *allometric heat-dissipation model*, which also revealed some differences in dispersal speed across locomotion modes. On average, flying animals can sustain potential dispersal speeds which are 100 times greater than those of running and swimming animals of equivalent body mass, while the potential dispersal speeds of swimming animals are marginally faster than those of running animals (Parameter *v*_0_ in Table 3). Flying animals’ higher potential dispersal speed, however, lead to the earlier saturation and subsequent decrease in their realised dispersal speed with increasing body mass (Fig 2).

**Table 3.** Posterior parameter estimates of the best-performing allometric locomotion model, the *allometric heat-dissipation model* (see Table 2 and S1 Table). Table entries correspond to the mean, standard deviation (SD), and 90% credible intervals of the model parameters’ posterior distributions (see Table 1 for a description of the model parameters).

| Parameter | Mean | SD | 5% | 95% |
| --- | --- | --- | --- | --- |
| $v_0$ (flying) | 109.16 | 9.37 | 95.00 | 125.06 |
| $v_0$ (running) | 0.99 | 0.07 | 0.88 | 1.11 |
| $v_0$ (swimming) | 1.39 | 0.10 | 1.24 | 1.56 |
| $k_\lambda$ | 0.0091 | 0.0013 | 0.0070 | 0.0113 |
| $c$ | 0.27 | 0.0078 | 0.25 | 0.28 |
| $d$ | 0.24 | 0.034 | 0.18 | 0.29 |

**Fig 2.**
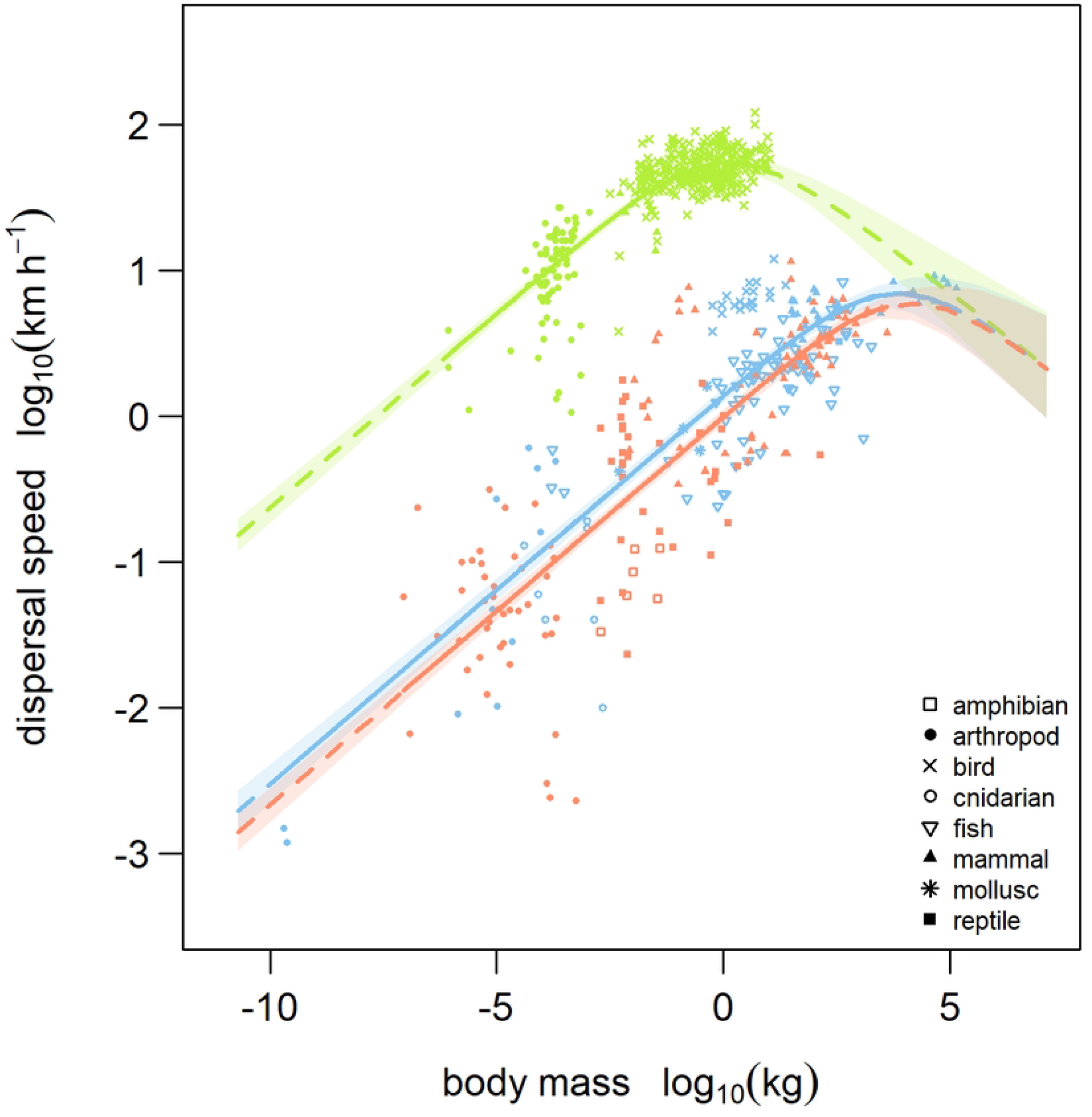
Realised dispersal speed as a function of body mass and locomotion mode as predicted by the *allometric heat-dissipation model*. Model-predicted mean values and 90% credible intervals are shown for flying (green), running (red), and swimming (blue) animals. The locomotion rate constant, *v*_0_, is fitted independently (i.e. no pooling) for each locomotion mode. Solid lines are predictions from the empirically observed range of body masses within each respective locomotion mode and dashed lines are predictions extrapolated beyond that range.

The *allometric heat-dissipation model* incorporates two allometric scaling exponents that characterise the body-mass constraints on (1) metabolic power input and locomotion efficiency (*c* in Table 1 and Table 3), and (2) heat-dissipation time (*d* in Table 1 and Table 3). The full model derivation (equations 2-6 and 20-27 in S1 Text) yields the expected ranges for these scaling exponents according to the underlying processes: The allometric scaling exponent which describes the initial increase in potential dispersal speed with increasing body mass, and thus, larger animals’ greater capacity to supply metabolic power combined with their higher locomotion efficiency, fell within the range of these theoretical expectations (expected: 0.01 < *c* < 0.*33*; observed: *c =* 0.2*7*, 90% CI 0.25 to 0.28). Similarly, the allometric scaling exponent that describes the body-mass dependence of heat-dissipation time, and thus, larger animals’ greater thermal inertia, also fell within the range of values derived from theoretical and empirical estimates of maximal heat-dissipation capacity (expected: 0.01 < *d* < 0.37; observed: *d =* 0.2*4*, 90% CI 0.18 to 0.29). Together, these results support the conclusion that these different allometric scaling processes jointly influence the realised dispersal speed.

An important question that remains is whether other variables might gain importance in explaining the deviations from our model predictions. The flying, running, and swimming animals included within our dataset cover a wide range of taxonomic groups (amphibians, arthropods, birds, cnidarians, fishes, mammals, molluscs, reptiles), thermoregulatory strategies (ectotherm, endotherm, mesotherm) and primary diets (carnivore, omnivore, herbivore), and these variables could contribute to explaining some of the variation in dispersal speed. Although our model did not account for these additional covariates, we found that they did not correspond to systematic deviations from our predicted values (S4-6 Figs). Since we tested our models based on a priori mechanistic derivations, we refrained from including these covariates in the statistical models.

## Discussion

We found a hump-shaped scaling relationship of dispersal speed with body mass across running, flying and swimming animals, which we explain using fundamental biophysical constraints on the supply, utilisation and dissipation of energy and heat, as incorporated within our *allometric heat-dissipation model*. This leads to two general insights about the fate of metabolic energy that goes towards the production of mechanical work and that which is dissipated as heat: First, despite possessing the metabolic potential to sustain higher dispersal speeds, the realised dispersal speeds of the largest animals are limited due to the risk of hyperthermia. Second, flying animals sustain a higher metabolic power input and higher dispersal speeds and, therefore, begin to limit their realised dispersal speeds at smaller body masses than running or swimming animals that travel more slowly. By jointly considering how allometric constraints shape metabolic demands as well as the dissipation of heat, we can provide generalised predictions of animal dispersal speed across different locomotion modes and ecosystem types when only the body mass is known.

Currently established models of animal locomotion typically produce power-law scaling relationships between locomotion speed and body mass by considering the dominance of a particular set of biophysical constraints on one critical aspect of the locomotion process; for example, biomechanical safety factors and the risk of physical injury [25,**Error! Reference source not found**.], dynamic similarity in locomotor mechanics and stride length [**Error! Reference source not found**.,**Error! Reference source not found**.], and the availability of metabolic power [34,48]. The scientific elegance of these biophysical models is that they relate a real-world phenomenon such as the speed of animal locomotion to the first principles of physics and morphology. Nevertheless, one of the limitations shared by most power-law models has been their limited ability to generalise predictions of animal locomotion speeds across a sufficiently wide range of body masses and across taxonomic groups that vary considerably in their body plan and mode of locomotion (e.g. [34,**Error! Reference source not found**.-**Error! Reference source not found**.**Error! Reference source not found**.]). Although such models describe how a particular biophysical constraint influences the utilisation of energy by the locomotory musculature, they do not take into account the considerable fraction of the total metabolic demand that is dissipated internally as heat. This leads to predictions such as those of the power-law model in our study (*the metabolic model*), which, by overestimating the sustained speeds of the largest animals, fails to capture the empirical data’s hump-shaped trends in log-log space. To address this, existing power-law models report values of the allometric scaling exponent which are inconsistent across taxonomic groups and different body mass ranges: One pattern which emerges across studies is the tendency for larger scaling exponents to be reported within groups of small-bodied animals such as arthropods [37,38,52], whereas large-bodied animals such as vertebrates tend to exhibit scaling relationships with smaller [21,**Error! Reference source not found**.,48], mass-independent [53,**Error! Reference source not found**.], or even negative scaling exponents [**Error! Reference source not found**.]. We have developed a biophysical model that reconciles these idiosyncrasies by incorporating both the allometry of locomotion efficiency (i.e. larger animals have a lower metabolic cost of locomotion) and the allometry of heat-dissipation capacity (i.e. larger animals require more time to dissipate metabolic heat) which, together, explain the initial increase, saturation, and inevitable decrease in realised dispersal speed with increasing body mass across each locomotion mode. The hump-shaped scaling relationship predicted by the *allometric heat-dissipation model* captures these trends in dispersal speed across the full range of animal body masses in our empirical dataset (from 2.00 × 10^−10^ to 1*4*0,000 *kg*), thereby setting universal limits to the sustained speed of any dispersing animal that relies on cyclical muscle contractions to fuel the performance of mechanical work.

The overall hump-shaped trend in the allometric scaling of dispersal speed is well-supported in our data by the dispersal speeds recorded among the world’s largest animals across various ecosystem types. For example, the median cruising speeds (± SD) of fin whales (8.64 ± 1.87 km/h, 74,000 kg) and blue whales (7.49 ± 1.66 km/h, 140,000 kg) reported by Gough et al. [**Error! Reference source not found**.] agree much more closely with the predictions of our *allometric heat-dissipation model* (fin whale: 5.85 km/h, 90% PI 1.67 to 21.72; blue whale: 5.37 km/h, 90% PI 1.45 to 19.67) than with those of the *metabolic model* (fin whale: 15.39 km/h, 90% PI 4.27 to 56.7; blue whale: 18.50 km/h, 90% PI 5.06 to 61.37). Moreover, the assumption that heat dissipation is a constraint to sustained locomotion also corresponds to some well-documented forms of behavioural thermoregulation. For instance, most migrating birds fly at high altitudes or at night in colder air, which, in turn reduces evaporative water losses [55,**Error! Reference source not found**.]. These behaviours are associated with a lower oxygen availability and an increased mechanical cost of flight at altitude [57,58] or the failure to settle at suitable stopover sites (night flights, [59]) and, therefore, only remain beneficial when heat dissipation is critical to sustained locomotion. There is also evidence across various taxonomic groups that the endogenous heat produced during locomotion can contribute to thermoregulatory requirements, for example, by allowing small mammals and birds to offsets a portion of the thermoregulatory cost associated with activity in colder climates [**Error! Reference source not found**.,**Error! Reference source not found**.] or by facilitating a form of intermittent endothermy among some the largest flying insects such as sphinx moths [62,**Error! Reference source not found**.]. Together, these examples illustrate the importance of metabolic heat production and dissipation for actively dispersing animals, which lies at the core of our new model of sustained speed. Interestingly, a similar hump-shaped relationship has also been shown to characterise the scaling of maximum speed with body mass by considering the combination of a finite time available for acceleration, which is limited due to the restricted availability of energy stored in the muscle cells to fuel anaerobic metabolism, together with body mass-constraints on animals’ energy storage capacity [40]. Surprisingly, this suggests that maximum speed and dispersal speed, although both hump-shaped in relation to body mass, could nevertheless be constrained by very different physiological processes that take precedence during short anaerobic bouts and sustained aerobic activity, respectively. This discrepancy highlights the importance of ecological context for understanding the processes that limit the performance of animals in different behavioural states.

Consistent with previous models [37,**Error! Reference source not found**.], we show that sustained speed initially scales with an allometric exponent close to 0.26. This allometric scaling exponent emerges from the product of the allometries of maximal aerobic metabolism (scaling with an exponent between 0.80 and 0.97 [65-**Error! Reference source not found**.]) and maximum locomotion efficiency (scaling with an exponent of approximately −0.67 [**Error! Reference source not found**.-**Error! Reference source not found**.**Error! Reference source not found**.]). While our statistical approach does not allow us to disentangle the relative contribution of these two interacting processes, the expected value of the exponent (between 0.13 and 0.30) agrees well with the empirically determined value of 0.26, supporting our model assumptions. This allometric scaling relationship holds until it reaches a saturation phase that characterises the maximum realised dispersal speeds within each locomotion mode. We found that this saturation phase and the subsequent decrease in realised dispersal speeds with increasing body mass occurred sooner in flying animals, between 0.1 kg (e.g. common starling) and 1 kg (e.g. herring gull), than in running or swimming animals (between 1,000 and 10,000 kg, e.g. Elephant or Orca). This discrepancy may be explained by the fact that flying animals sustain higher rates of aerobic metabolism during locomotion than running and swimming animals [58,**Error! Reference source not found**.] and utilise muscles that operate with lower mechanical efficiencies [70,**Error! Reference source not found**.]. Consequently, they encounter the limits of their allometric heat-dissipation capacity at a smaller body mass. Following this initial saturation with increasing body mass, realised dispersal speed ultimately tends towards an exponent of −0.24 across all locomotion modes. This corresponds well with our model assumptions for the allometric scaling of heat-dissipation time (equations 20-27 in S1 Text) as well as with the observation many large flying birds sustain flight speeds that are lower than those that would maximise their migration range or aerobic efficiency [**Error! Reference source not found**.]. In addition to the general similarity in the hump-shaped scaling relationship across locomotion modes, our study also quantifies important differences between running, flying and swimming animals.

Overall, flying animals are able to sustain much greater speeds than running or swimming animals of equivalent body mass. This is driven by an almost 100-fold larger value of their locomotion rate constant, *v*_0_, which encompasses the mass-independent interaction between the rate of aerobic metabolism and locomotion efficiency. Our model does not assume that metabolic power input and locomotion efficiency vary independently of one another [70], but rather, that there are systematic differences among flying, running and swimming animals in their maximal capacities to (i) supply their muscles with metabolic energy via aerobic pathways [58,**Error! Reference source not found**.] and (ii) utilise this energy efficiently during locomotion [**Error! Reference source not found**.- **Error! Reference source not found**.**Error! Reference source not found**.]. Overall, our *allometric heat-dissipation model* not only predicts the hump-shaped relationship between realised dispersal speed and body mass but also provides an explanation for the differences in the shape of this scaling relationship between locomotion modes.

We have derived the *allometric heat-dissipation model* from physical first principles based on the core mechanistic components of (1) metabolic energy supply, (2) the metabolic cost of locomotion, and (3) heat-dissipation capacity. The fit of our model to empirical data yielded a general parametrisation that will allow future predictions of animal dispersal rates based on body mass and locomotion mode as the only species traits. The simplicity of the model structure and generality of its applicability come at the expense of excluding additional factors that may affect the speed of locomotion without universally affecting any of the three core mechanistic model components: This includes, for example, phylogenetic history or morphological constraints (e.g. limb length, hovering versus forward flapping flight). While the inclusion of these covariates could improve the prediction of dispersal speeds for specific groups of animals, it would come at the expense of generality in model predictions across all locomotion types. For example, species’ limb length could be added to better predict the dispersal speed of running animals [32,73], whereas this component would be less meaningful when considering the effects of lift and drag on flying and swimming animals, respectively. Similarly, we chose to exclude information on phylogenetic relatedness because the biophysical principles included in our model are universal and deeply rooted in evolutionary history (e.g. total mitochondrial volume, protein temperature-dependence) and should apply, therefore, to all taxonomic groups. From a philosophical perspective, the use of phylogenetic covariates to improve the model fit would defeat the purpose of developing a universal model based on biophysical first principles. Therefore, we have currently limited our approach to biophysical processes that can be generalised across all modes of locomotion and across taxonomic groups. Nevertheless, there is scope for additional environmental or morphological factors that affect any of the three core mechanistic model components to be included as extensions of our model. The importance of the aforementioned constraints can be identified by analysing deviations from our model predictions: For instance, Arctic wolves (*Canis lupus*) on Ellesemere Island [74] - the most northerly island within the Arctic Archipelago - travel 3.3 km/h faster than the upper bound of the 90% prediction interval of our *allometric heat-dissipation model*. We anticipate that they are able to sustain such high speeds over distances of 2-4 km while returning to their summer dens because the ambient temperatures that they experience rarely exceed −1.0 degrees C, thus enabling a more rapid dissipation of the heat generated by muscular thermogenesis. This illustrates an important effect of low ambient temperature on reducing the time required for heat dissipation which, in turn, increases the thermoregulatory capacity (of large animals) to sustain high dispersal speeds. In its current form, our *allometric heat-dissipation model* includes the simplifying assumption that core body temperature increases with distance travelled without specifically considering the temperature of the body or that of the ambient environment. Incorporating temperature data into our model could therefore further elucidate additional physiological limits to animals’ capacities for sustained locomotion. We also found that morphological defence traits such as shell of the Galapagos giant tortoise (*Chelonoidis niger*) - the largest extant terrestrial ectotherm – coincides with sustained speeds that are considerably slower than the lower bound of our model’s 90% prediction interval. We propose that the incredibly slow walking speeds of Galapagos giant tortoises arise because of their shell’s limited capacity to dissipate endogenously produced heat rather than because of its weight [**Error! Reference source not found**.]. This suggests an interesting trade-off between local persistence through the defence against natural enemies and the capacity to disperse to distant but (potentially) predator-free environments.

Moreover, the evolution of morphological adaptations that facilitate heat dissipation (e.g. counter-current systems [**Error! Reference source not found**.], the avian bill [**Error! Reference source not found**.]) may have important implications for the dispersal capacities of animals in ancient and contemporary landscapes. Together, these examples illustrate how additional characteristics of a species’ morphology and its thermal environment could be incorporated into our *allometric heat-dissipation model*. The model, thereby, retains its generality across a wide range of taxonomic groups and locomotion modes by including the quantitative responses of model components such as heat-dissipation capacity to these characteristics.

## Conclusions

Animal dispersal plays a critical role in shaping ecological dynamics at larger spatial scales [78-**Error! Reference source not found**.**Error! Reference source not found**.]. Realistic models of landscape-scale biodiversity dynamics must incorporate large numbers of species whose dispersal rates can be predicted only on the basis of easily quantifiable traits such as body mass and locomotion mode. Therefore, allometric locomotion models such as our *allometric heat-dissipation model* could provide the modular “building blocks” of dynamic meta-community or meta-food web models. This has a great potential to reveal the interplay between spatial processes and species interactions in driving biodiversity patterns across spatial scales [8,**Error! Reference source not found**.,81,**Error! Reference source not found**.]. In contrast to existing power-law models [21,34-**Error! Reference source not found**.**Error! Reference source not found**.38], our *allometric heat-dissipation model* accurately predicts that sustained dispersal speed follows a hump-shaped relationship with body mass. With respect to the spatial processes linking local communities to one another within metacommunities such as forest- or island archipelagos, the identification of the novel link between metabolic heat-dissipation constraints and animal dispersal capacity implies that large animal species may be more susceptible to the effects landscape fragmentation than previously anticipated [17,**Error! Reference source not found**.]. Specifically, the larger total metabolic energy expenditure associated with increasing animal body mass needs to be balanced by an increased ability to track spatial resource dynamics at the landscape scale. Our results suggest that, as animal body mass increases beyond the critical threshold defined by the saturation phase of our locomotion model, further increases in total metabolic demand will coincide with a decreasing dispersal capacity.

Consequently, the high extinction risk observed among large animals, especially terrestrial herbivores and reptiles [**Error! Reference source not found**.], may also be driven by their inability to balance their metabolic demands by efficiently locating resources within patchy landscapes [85,**Error! Reference source not found**.]. Our *allometric heat-dissipation model* helps to reconcile animal dispersal theory with empirical biodiversity patterns and underpins the novel call to protect large animals from the dire consequences of landscape fragmentation.

## Materials and Methods

### The dataset

We synthesised a dataset containing empirical estimates of sustained dispersal speed for flying, running, and swimming animals by searching ISI Web of Science and Google scholar for published studies using the following keywords: (optimal OR cruising OR travel OR routine OR dispersal OR sustained) AND (velocity* OR speed* OR motilit*) AND (run* OR walk* OR terrestrial OR flight* OR fly* OR swim*). We also searched the bibliographies of relevant publications for additional data sources.

We included data from field and laboratory studies that reported mean or median speeds of individual animals or groups of animals maintaining sustained and directed movements within an unrestrained setting. This precluded the use of movement data obtained from treadmills, flight mills, swim tunnels, wind tunnels, as well as from animals who were stimulated to move by an observer. Studies estimated dispersal speeds either directly (i.e. instantaneously) through visual observations, video recordings, radar, and animal-attached devices or indirectly from higher-frequency (sampling interval < 30 minutes) telemetry data. For flying animals, we only considered flight speeds during powered (i.e. thrust generated by flapping) flight. When individual- or species-level body mass was not provided, we referred to secondary literature sources to assign the average adult body mass of the species (e.g. [**Error! Reference source not found**.]). In cases where only body length was given, we used published allometric equations to estimate the wet body mass (e.g. [**Error! Reference source not found**.]). For studies that reported individual-level data, we aggregated data to the species level by calculating the unweighted geometric mean of individual dispersal speeds and, where available, individual body masses. This resulted in a dataset (S1 Data) that featured 697 estimates of mean or median dispersal speed taken from 169 studies across a pool of 531 species from various taxonomic groups (amphibians, arthropods, cnidarians, birds, fishes, mammals, molluscs, reptiles) that spanned 15 orders of magnitude in body mass (from 2.00 × 10^−10^ to 140,000 kg) and 5 orders of magnitude in dispersal speed (from 1.1*9* × 10^−3^ to 121 km/h)

### Model specification

We used Bayesian parameter estimation to evaluate the relationship between body mass and dispersal speed. Bayesian models are comprised of three components: (i) a stochastic data model that links model predictions to the observed data; (ii) a deterministic process model that describes each of our mechanistic hypotheses; (iii) a parameter model that includes prior assumptions about the parameter values.

#### (i) Data model

For the data model, we assumed a Gaussian likelihood for the log_10_-transformed dispersal speed *v*_*i*_ (in km/h):

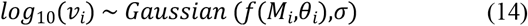

where the process model, *f*(·), predicts expected dispersal speed from the observed body mass, *M* (in kg) the vector of observation-level parameters, *θ*_*i*_ *= {c,d,v*_0_,*k*_0_,*k*_*λ*_*}*, and the standard deviation, *σ*, between model predictions and the observed data.

#### (ii) Process model

We considered three alternative process models of varying complexity which corresponded to our three alternative hypotheses (Table 1 step 8 and Fig 1D) about the form of the allometric scaling relationship for realised dispersal speeds. Each process model was reformulated in log_10_-linear form. We included locomotion mode as a categorical covariate by estimating the locomotion rate constant (parameter *v*_0_) independently (i.e. no pooling) for flying, running, and swimming animals; the normalisation constants associated with heat-dissipation time (*k*_0_ or *k*_*λ*_) did not vary among locomotion modes (i.e. complete pooling).

#### (iii) Parameter model

We specified weakly informative prior distributions in the parameter model. The observation-level parameters for the allometric scaling exponents *c* and *d* were assumed to have independent normal priors; observation-level parameters for the normalisation constants *v*_0_, *k*_0_, and *k*_*λ*_ were assigned independent gamma distribution priors. We assumed a half-Cauchy prior distribution for the observation-level variances.

### Model selection and inference

Model selection and inference included the evaluation of the alternative allometric process model formulations. We fitted each model using a No-U-Turn Hamiltonian Monte Carlo Sampler (NUTS-HMC) in Stan via the rstan package in R 4.0.2 by employing three parallel NUTS-HMC chains with an adaptation phase of 1,500 iterations and a sampling phase of 3,000 iterations each. This yielded a sum of 9,000 samples of the posterior distribution for each model. A visual assessment of the trace- and density plots confirmed that the NUTS-HMC chains had mixed adequately; Gelman-Rubin statistics ≤ 1.01 and high (>1000) effective sample sizes verified convergence [89]. We compared the out-of-sample prediction accuracies of each model by calculating the LOOIC (leave-one-out information criterion) from the expected log predictive densities (ELPD) using the log-likelihood values of the posterior samples (R package loo). We used a difference in LOOIC (ΔLOOIC) which was larger than at least two times the estimated standard error of the difference (SE ΔLOOIC) to distinguish among competing models [**Error! Reference source not found**.]. Samples from the posterior distribution were used to characterise the distribution of parameter values and to estimate model uncertainty by reporting central and 90% credible intervals (CIs) for parameter estimates as well central and 90% prediction intervals (PIs) from model-based predictions of realised dispersal speed.

## Acknowledgements

A. Dyer, E. Berti, M.R. Hirt, B. Rosenbaum and U. Brose acknowledge the support of the German Centre for Integrative Biodiversity Research Halle-Jena-Leipzig, funded by the German Research Foundation: FZT 118, 202548816 and the research group DynaCom (spatial community ecology in highly dynamic landscapes: from island biogeography to metaecosystems) funded by the German Research Foundation (DFG, FOR 2716).

## Author contributions

Conceptualization: Alexander Dyer, Myriam Hirt, Ulrich Brose

Data curation: Alexander Dyer, Myriam Hirt

Formal analysis: Alexander Dyer, Emilio Berti, Benjamin Rosenbaum

Methodology: Alexander Dyer, Emilio Berti, Myriam Hirt, Ulrich Brose

Visualisation: Alexander Dyer, Emilio Berti, Myriam Hirt, Ulrich Brose

Writing - original draft preparation: Alexander Dyer, Myriam Hirt, Ulrich Brose

Writing - review & editing: Alexander Dyer, Emilio Berti, Myriam Hirt, Benjamin Rosenbaum, Ulrich Brose

## Supporting information

**S1 Fig. Predictions from the metabolic model for realised dispersal speed as a function of body mass**. Model-predicted mean values and 90% credible intervals are shown for flying (green), running (red), and swimming (blue) animals. The locomotion rate constant, *v*_0_, is fitted independently (i.e. no pooling) for each locomotion mode. Solid lines are predictions from the empirically observed range of body masses and dispersal speeds within each respective locomotion mode and dashed lines are predictions extrapolated beyond that range.

(S1_Fig.TIFF)

**S2 Fig. Predictions from the constant heat-dissipation model predictions for realised dispersal speed as a function of body mass**. Model-predicted mean values and 90% credible intervals are shown for flying (green), running (red), and swimming (blue) animals. The locomotion rate constant, *v*_0_, is fitted independently (i.e. no pooling) for each locomotion mode. Solid lines are predictions from the empirically observed range of body masses within each respective locomotion mode and dashed lines are predictions extrapolated beyond that range.

(S2_Fig.TIFF)

**S3 Fig. Predictions from the allometric heat-dissipation model for realised dispersal speed as a function of body mass with the allometric scaling exponent *c* fitted independently (i**.**e. no pooling) for each locomotion mode**. Model-predicted mean values and 90% credible intervals are shown for flying (green), running (red), and swimming (blue) animals. The locomotion rate constant, *v*_0_, is fitted independently (i.e. no pooling) for each locomotion mode. Solid lines are predictions from the empirically observed range of body masses within each respective locomotion mode and dashed lines are predictions extrapolated beyond that range.

(S3_Fig.TIFF)

**S4 Fig. Effect of taxonomic group on the realised dispersal speed of animals**. Residual deviations from the mean dispersal speeds predicted by the allometric heat-dissipation model (Fig 2). Box plots show median deviations (horizontal line), 25th and 75th percentiles (boxes), extreme values (whiskers).

(S4_Fig.TIFF)

**S5 Fig. Effect of thermoregulatory strategy on the realised dispersal speed of animals**. Residual deviations from the mean dispersal speeds predicted by the allometric heat-dissipation model (Fig 2). Box plots show median deviations (horizontal line), 25th and 75th percentiles (boxes), extreme values (whiskers).

(S5_Fig.TIFF)

**S6 Fig. Effect of diet on the realised dispersal speed of animals**. Residual deviations from the mean dispersal speeds predicted by the allometric heat-dissipation model (Fig 2). Box plots show median deviations (horizontal line), 25th and 75th percentiles (boxes), extreme values (whiskers).

(S6_Fig.TIFF)

**S1 Table.** Comparison of six alternative allometric locomotion models that predict the realised dispersal speeds of animals as a function of their body mass and locomotion mode. The more complex versions of each of the allometric locomotion models featured in Table 1 also allow for variation in the allometric scaling exponent *c* among flying, running and swimming animals. LOOIC values are presented together with the difference in LOOIC value relative to the most parsimonious model (ΔLOOIC = 0.0) and the estimated standard error of the difference (SE ΔLOOIC). LOOIC represents the expected log pointwise-predictive densities (ELPD) converted to the deviance scale. The difference in LOOIC (ΔLOOIC) between the two best-fitting models, the *allometric heat-dissipation model* with shared versus variable slope, is within two standard errors of the difference (SE ΔELPD) indicating that there is no distinguishable difference in their predictive performance.

| Model | Description | LOOIC | $\Delta\text{LOOIC}$ | $\text{SE } \Delta\text{LOOIC}$ |
| --- | --- | --- | --- | --- |
| Metabolic (shared slope) | Power law: only $v_0$ varies with locomotion mode | 465.0 | 64.8 | 14.2 |
| Metabolic (variable slope) | Power law: $v_0$ and $c$ vary with locomotion mode | 465.0 | 64.8 | 14.2 |
| Constant heat-dissipation (shared slope) | Saturating: only $v_0$ varies with locomotion mode | 427.6 | 27.5 | 8.1 |
| Constant heat-dissipation (variable slope) | Saturating: $v_0$ and $c$ vary with locomotion mode | 414.2 | 14.1 | 5.5 |
| Allometric heat-dissipation (shared slope) | Hump-shaped: only $v_0$ varies with locomotion mode | 402.2 | 2.0 | 5.3 |
| Allometric heat-dissipation (variable slope) | Hump-shaped: $v_0$ and $c$ vary with locomotion mode | 400.1 | 0.0 | 0.0 |

**S1 Text. Model derivation**.

(S1_Text.DOCX)

**S1 Data. Dispersal speed and body mass data**.

(S1_Data.CSV)

## Notes

### Competing Interest Statement

The authors have declared no competing interest.

